# Differential urine proteome analysis of a bovine IRBP induced uveitis rat model by discovery and parallel reaction monitoring proteomics

**DOI:** 10.1101/782045

**Authors:** Weiwei Qin, Lujun Li, Ting Wang, Youhe Gao

## Abstract

Uveitis, a group of intraocular inflammatory diseases, is one of the major causes of severe visual impairment among working age population. Urine is a promising resource for biomarker research, which could sensitively reflect the changes of the body. This study was designed to explore urinary protein biomarkers for diagnosis and/or monitoring of uveitis. Experimental autoimmune uveitis (EAU) rat model induced by bovine interphotoreceptor retinoid-binding protein (IRBP) was used to mimic the uveitis. In the discovery phase, urine samples from EAU and control rats were analyzed by data independent acquisition (DIA) approach combined with high-resolution mass spectrometry. Overall, 704 high confidential proteins were identified, of which 100 were differentially expressed (37, 33, 37, and 44 on day 5, 8, 12, and 16, respectively, after bovine IRBP immunization) (1.5-fold change, P<0.05). Gene Ontology analysis of the dysregulated proteins showed that chronic inflammatory response, neutrophil aggregation and immune system processes were significantly enriched. Finally, parallel reaction monitoring (PRM) approach was used for further validation. A total of 12 urinary proteins (MMP8, NGAL, HPT, UROM, RISC, A1AG, TTHY, KNT1, C9, PTER, CBG, and FUCA1) changed significantly, even when there is no clinical manifestations and histopathological ocular damages in the EAU rats. Our findings represent the first step towards urinary protein diagnostic biomarkers for uveitis.

**Biological Significance:** This is the first study investigating urinary protein candidate biomarkers for uveitis using data independent acquisition (DIA) combined with parallel reaction monitoring (PRM) in experimental autoimmune uveitis (EAU) rats. The results revealed that urine is a promising resource for early diagnostics of uveitis. Further research including clinical urine samples is needed to determine the sensitivity and specificity of these candidate biomarkers for uveitis.

## 1. Introduction

Uveitis is a group of intraocular inflammatory diseases which mainly involve uvea, the blood-vessel rich pigmented middle layer of the wall of the eye. Uveitis carries a high risk of vision loss, and usually affects people aged 20-50 years [1–3]. In developing countries, approximately 5-25% of irreversible blindness is caused by uveitis and its complications [4, 5]. In addition to the infectious agents, autoimmune reaction is one of the main causes of uveitis. As for autoimmune uveitis, corticosteroids and immunomodulatory agents are the mainstay of treatment [6]. However, there are some uveitis patients who fail to respond to the treatment. And it’s common for the recurrence to occur during the tapering phase of corticosteroids. In addition, the long-term treatments may have several intraocular and systemic side effects, such as high intraocular pressure, sterile endophthalmitis, and nephrotoxicity [7, 8].

The diagnosis and differential diagnosis of uveitis remains challenging even to experienced specialists [9, 10]. On one hand, the profound clinical overlap between uveitis entities induced by different etiologies; on the other, the limited availability of clinical specimen and reliability/accuracy of currently available tests. Timely diagnosis of non-autoimmune uveitis, in particular infectious uveitis and masquerade syndrome, can be difficult. Therefore, identifying simple and accurate biomarkers for diagnosis and/or disease monitoring are of great clinical significance. If diagnosed earlier, effective measures may be used to prevent vision loss.

Urine is an attractive resource for biomarker research that has been underutilized. Without strict homeostatic regulation, urine can sensitively reflect changes in the body at early stage [11, 12]. Mass spectrometry-based proteomics has dramatically improved and emerged as a prominent tool in the field of biomarker studies. Many candidate biomarkers of uveitis have been described primarily in blood, tear, and aqueous humor [13–16]. Urinary proteomic studies have identified many candidate biomarkers for autoimmune inflammatory disease, such as, rheumatoid arthritis, autoimmune myocarditis, inflammatory bowel disease [17, 18]. However, there have been limited studies of protein biomarker in urine for uveitis, compared with the wide applications for other diseases.

Experimental autoimmune uveitis (EAU) is T cell-mediated autoimmune disease that targets the neural retina and related tissues [19, 20]. It is an induced noninfectious uveitis, as opposed to spontaneous, autoimmune disease model. The hallmarks of EAU are onset of ocular inflammation, disruption of the retinal architecture, and partial to complete destruction of the photoreceptor cell layer [21]. It shares many common features in clinical and histological aspects with human uveitis [22]. Urine proteome is affected by multiple factors, such as age, diet, exercise, gender, medication, and daily rhythms [23]. Animal models can be used to minimize the impact of many uncertain factors by establishing a direct relationship between a disease and corresponding changes in urine [24].

This study aimed to identify the potential urinary protein biomarkers related to uveitis by using the bovine IRBP induced uveitis rat model. The experiment was conducted in two phases. In the discovery phase, data-independent acquisition (DIA) approach was used to profile the proteome of urine from EAU rats, and compared them with the controls. In the validation phase, the differentially expressed proteins were validated by parallel reaction monitoring (PRM) targeted quantitative analysis using a quadrupole-orbitrap mass spectrometer.

## 2. Materials & Methods

### 2.1 Animals and Uveitis Induction

Fifty male Lewis rats (160–180 g) were purchased from Charles River China (Beijing, China). All animals were maintained with a standard laboratory diet under controlled indoor temperature (21±2°C), humidity (65–70%) and 12 h light#dark cycle conditions. The animal experiments were reviewed and approved by the Peking Union Medical College (Approved ID: ACUC-A02-2014-008) and performed in accordance with the guidelines for animal research.

Lewis rats were randomly divided into two groups: the control group (n = 25) and the EAU group (n = 25). EAU was induced according to the procedure described previously [21]. Briefly, rats in the EAU group were immunized with 30 μg bovine IRBP peptide R16 (ADGSSWEGVGVVPDV, BGI, Beijing, China), emulsified 1:1 in complete Freund’s adjuvant (Sigma-Aldrich, St. Louis, MO, USA). Rats in the control group were given a subcutaneous injection of same volume of normal saline.

The experiment was conducted in two phases, for the details in figure 1. For the discovery phase, differential urinary proteins were identified by DIA label free quantification, in forty independent samples from the control group (8 samples) and the EAU group on day 5, 8, 12 and 16 (8 samples per time point); and for the validation phase, urine samples obtained from the 32 remaining urine samples (8 from the control group and the other from the EAU group on day 5, 8 and 12, 8 samples per time point) by PRM targeted quantification.

**Figure 1.**
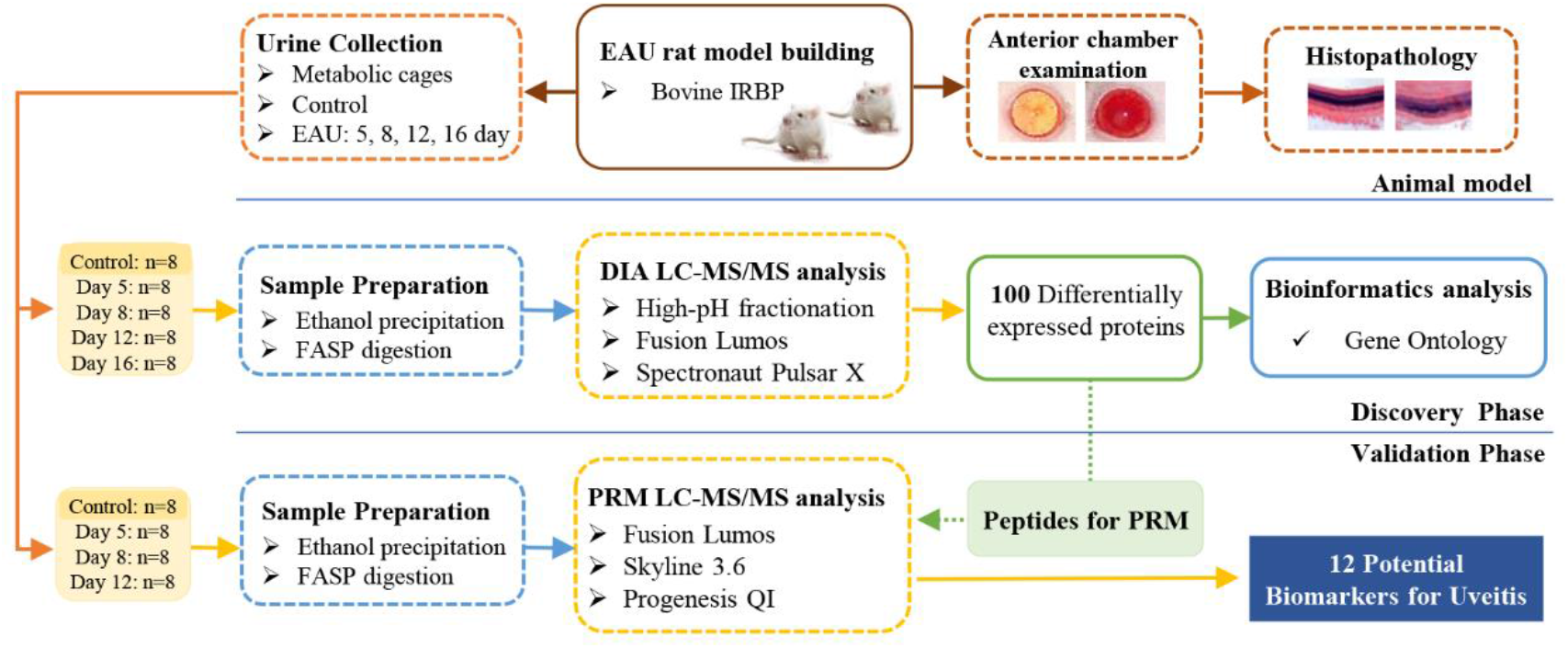
Workflow of the study of urine proteome changes in experimental autoimmune uveitis (EAU) rats.

### 2.2 Histological Analysis of EAU

For histopathology, three rats in the experimental group and three rats in the control group were randomly sacrificed at 8, 12 and 16 days after immunization injection. The eye balls were harvested and then quickly fixed in 10% neutral-buffered formalin. The formalin-fixed tissues were embedded in paraffin, sectioned (4 mm) and stained with hematoxylin and eosin (H&E) to reveal histopathological lesions.

### 2.3 Urine Collection and Sample Preparation

Urine samples were collected from the control and experimental groups at days 5, 8, 12 and 16 after immunization injection. Rats were individually placed in the metabolic cages for six hours. During urine collection, no food was provided for rats to avoid urine contamination. After collection, urine samples were centrifuged at 2 000 g for 30 min at 4°C immediately, and then stored at −80 °C.

Urinary protein extraction: urine samples were centrifuged at 12 000g for 30 min at 4 °C. Six volumes of pre-chilled ethanol were added after removing the pellets, and the samples were precipitated at 4°C overnight. Then, lysis buffer (8 mol/L urea, 2 mol/L thiourea, 50 mmol/L Tris, and 25 mmol/L DTT) was used to re-dissolve the pellets. The protein concentration of each sample was measured by the Bradford protein assay. Tryptic digestion: the proteins were digested with trypsin (Promega, USA) using filter-aided sample preparation methods[25]. Briefly, 100 μg of the protein sample was loaded on the 10-kD filter unit (Pall, USA). The protein solution was reduced with 4.5 mM DTT for 1 h at 37°C and then alkylated with 10 mM of indoleacetic acid for 30 min at room temperature in the dark. The proteins were digested with trypsin (enzyme-to-protein ratio of 1:50) for 14 h at 37°C. The peptides were desalted on Oasis HLB cartridges (Waters, USA) and lyophilized for trap column fractionation and LC-MS/MS analysis.

### 2.4 Spin Column Separation

To generated a spectral library for DIA analysis, pooled peptides sample from all samples were fractionated using a high-pH reversed-phase peptide fractionation kit (Thermo Pierce, USA) according to the manufacturer’s instructions. Briefly, 60 ug pooled peptides sample was loaded onto the spin column. A step gradient of increasing acetonitrile concentrations was applied to the column to elute bound peptides. Ten different fractions were collected by centrifugation, including flow-through fraction, wash fraction and eight step gradient sample fractions (5, 7.5, 10, 12.5, 15, 17.5, 20 and 50% Acetonitrile). Fractionated samples were dried completely and resuspended in 20 μl of 0.1% formic acid. Three microliters of each of the fractions was loaded for LC-DDA-MS/MS analysis.

### 2.5 LC-MS/MS Setup for DDA and DIA

An Orbitrap Fusion Lumos Tribrid mass spectrometer (Thermo Scientific, Germany) was coupled with an EASY-nLC 1000 HPLC system (Thermo Scientific, Germany). For both DDA-MS and DIA-MS mode, the same LC settings were used for retention time stability. The digested peptides were dissolved in 0.1% formic acid and loaded on a trap column (75 μm × 2 cm, 3 μm, C18, 100 A°). The eluent was transferred to a reversed-phase analytical column (50 μm× 250 mm, 2 μm, C18, 100 A°). The eluted gradient was 5–30% buffer B (0.1% formic acid in 99.9% acetonitrile; flow rate of 0.4 μl/min) for 60 min. To enable fully automated and sensitive signal processing, the calibration kit (iRT kit from Biognosys, Switzerland) was spiked at a concentration of 1:20 v/v in all samples. The iRT kit was spiked into the urinary peptides for spectral library generation. Also, before the real DIA runs, the iRT kit was also spiked into all urinary samples.

For the generation of the spectral library, ten fractions from spin column were analyzed in DDA-MS mode. The parameters were set as follows: the full scan was acquired from 350 to 1 550 m/z at 60 000, the cycle time was set to 3 secs (top speed mode); the auto gain control (AGC) was set to 1e6; and the maximum injection time was set to 50 ms. MS/MS scans were acquired in the Orbitrap at a resolution of 15,000 with an isolation window of 2 Da and collision energy at 32% (HCD); the AGC target was set to 5e4, and the maximum injection time was 30 ms.

For the DIA-MS method, forty individual samples were analyzed in DIA mode. For MS acquisition, the variable isolation window DIA method with 26 windows was developed (Table S1). The full scan was set at a resolution of 60,000 over an m/z range of 350 to 1,200, followed by DIA scans with a resolution of 30,000, HCD collision energy of 32%, AGC target of 1e6 and maximal injection time of 50 ms. A quality control DIA analysis of the pooled sample was inserted after every 8 urine samples were tested.

### 2.6 LC-MS/MS Setup for PRM

In the discovery phase, fifty-nine differential expressed urinary proteins were identified by label-free DIA proteomic method. All these proteins were evaluated by PRM-MS method in the remining thirty-two urine samples. LC-PRM-MS/MS data acquired on a Orbitrap Fusion Lumos Tribrid mass spectrometer (Thermo Scientific, Germany) coupled with an EASY-nLC 1200 HPLC system (Thermo Scientific, Germany).

For the generation of the PRM spectral library, pooled peptide samples were analyzed in DDA-MS mode for 6 times. The peptides were loaded on a reversed-phase trap column (75 μm × 2 cm, 3 μm, C18, 100 Å, Thermo Scientific, Germany), and then the eluent was transferred to a reversed-phase analytical column (50 μm × 150 mm, 2 μm, C18, 100 Å, Thermo Scientific, Germany). The eluted gradient was 5–35% buffer B (0.1% formic acid in 80% acetonitrile; flow rate 0.3 μl/min) for 120 min. The MS parameters were set as follows: the full scan was acquired from 350 to 1 550 m/z at 60 000, the cycle time was set to 3 secs (top speed mode); the auto gain control (AGC) was set to 1e6; and the maximum injection time was set to 50 ms. MS/MS scans were acquired in the Orbitrap at a resolution of 30 000 with an isolation window of 1.6 Da and collision energy at 30% (HCD); the AGC target was set to 5e4, and the maximum injection time was 60 ms.

For the PRM-MS method, thirty-two individual samples were analyzed in PRM mode. Finally, 150 peptides were scheduled and the retention time (RT) segment was set to 8 min for each targeted peptide (Table S2). The normalized collision energy was fixed to 30%, and a quadrupole isolation window of 1.6 Da. The other parameters just as the same as described in the last paragraph.

### 2.7 The Label-free DIA Quantification Analysis

To generate the spectral library, the ten fractions’ raw data files acquired by the DDA mode were processed using Proteome Discoverer (version 2.3; Thermo Scientific, Germany) with SEQUEST HT against the SwissProt ratus database (released in May 2019, containing 8086 sequences) appended with the iRT peptides sequences. The Search parameters consisted of the parent ion mass tolerance, 10 ppm; fragment ion mass tolerance, 0.02 Da; fixed modifications, carbamidomethylated cysteine (+58.00 Da); and variable modifications, oxidized methionine (+15.995 Da) and deamidated glutamine and asparagine (+0.984 Da). Other settings included the default parameters. The applied false discovery rate (FDR) cutoff was 0.01 at the protein level. The results were then imported to Spectronaut™ Pulsar (Biognosys, Switzerland) software to generate the spectral library [26].

The DIA-MS raw files were imported to Spectronaut Pulsar with the default settings. In brief, a dynamic window for the XIC extraction window and a non-linear iRT calibration strategy were used. Mass calibration was set to local mass calibration. Cross-run normalization was enabled to correct for systematic variance in the LC-MS performance, and a local normalization strategy was used [27]. Protein inference, which gave rise to the protein groups, was performed on the principle of parsimony using the ID picker algorithm as implemented in Spectronaut Pulsar [28]. All results were filtered by a Q value cutoff of 0.01 (corresponding to an FDR of 1%). Peptide intensity was calculated by summing the peak areas of their respective fragment ions for MS2. The significance criteria for a T-test was a p value <0.05. A minimum of two peptides matched to a protein and a fold change >1.5 were used as the criteria for the identification of differentially expressed proteins.

### 2.8 The PRM-MS Quantification Analysis

Skyline (Version 3.6.1 10279) [29]was used to build the spectrum library and filter peptides for PRM analysis. For each targeted protein, 2-6 associated peptides were selected using the following rules: (i) identified in the untargeted analysis with q value <1%, (ii) completely digested by trypsin, (iii) contained 8–18 amino acid residues, (iv) the first 25 amino acids at the N-terminus of proteins were excluded, and (v) fixed carbamidomethylation of cysteine. Prior to individual sample analysis, pooled peptide samples were subjected to PRM experiments to refine the target list. Finally, forty-four proteins with 150 peptides (Table S2) were finally scheduled. The RT segment was set to 8 min for each targeted peptide with its expected RT in the center based on the pooled sample analysis. The technical reproducibility of the PRM assay was assessed, and the results showed that among the targeted peptides, 134 peptides have abundance CV values that are less than 30% (Fig. S2).

All of the PRM-MS data were processed with Skyline. By comparing the same peptide across runs, the RT location and integration boundaries were adjusted manually to exclude interfering regions. Each protein’s intensity was quantitated using the summation of intensities from its corresponding transitions. Transition settings: precursor charges +2, +3; ion charge +1; ion type b, y, p; product ions from ion 3 to last ion −1; auto-select all matching transitions; ion match tolerance 0.02 m/z; pick 6 most intense product ions. The details of the transition are listed in supporting Table S2. Prior to the statistical analysis, the quantitated protein intensities were normalized by the summed intensity. The differential proteins were selected using one-way ANOVA. Significance was accepted at a p-value of less than 0.05.

### 2.9 Bioinformatics analysis

Bioinformatics analysis was carried out in order to better study the biological function of the dysregulated proteins. GO analysis was performed on the urinary differentially altered proteins identified at the discovery phase (http://www.geneontology.org/) [30, 31]. In this study, significant GO enrichment was defined as P<0.05.

## 3. Results & Discussion

### 3.1 Clinical manifestations and histopathological ocular damage in EAU eyes

Anterior chamber examination and histopathology analysis were performed to evaluate the progression of EAU in the Lewis rats (Fig. 2). On day 8, there was no difference between the EAU and the control rats. Anterior chamber examination showed translucent appearance, the pupil and iris blood vessels were clearly visible and the vessels were not congested (Fig.2A). HE staining showed: the retina was in ordered retinal layers (Fig.2B). On day 12, it showed obvious signs of uveitis compared with control rats. The eyeball appeared larger due to swelling and proptosis, red reflex was absent and pupil was obscured (Fig.2A). HE staining showed: the retinal architecture was disorganized, massive inflammatory cell infiltrated throughout the retina and choroid, and photoreceptor cell was damaged (Fig. 2B). On day 16, it showed mild vitritis and retinal folds (Fig. 2).

**Figure 2.**
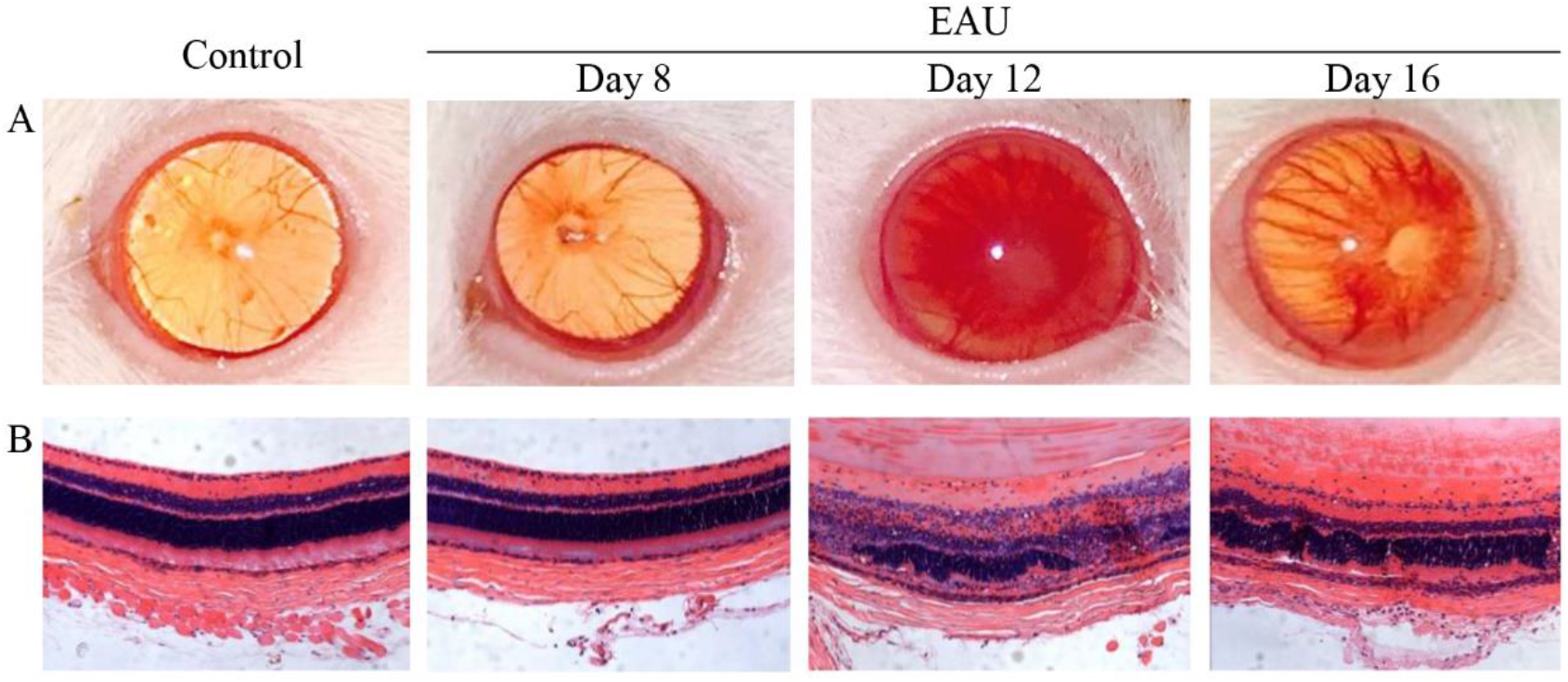
Clinical manifestations and histopathological ocular damage in EAU eyes. A: Clinical appearance by anterior chamber examination. B: Hematoxylin and eosin stain (HE) of retinal tissue (20×). Control: Normal control rat eye; Day 8: 8 days after the bovine IRBP immunization; Day 12: 12 days after the bovine IRBP immunization; Day 16: 16 days after the bovine IRBP immunization.

These clinical manifestations and pathological changes revealed the success of the bovine IRBP induced uveitis modeling. And EAU severity peaked on day 12 after the immunization injection.

### 3.2 Urine proteome changes between the EAU and control rats

To preliminary investigate how the urine proteome changes with EAU progression, forty urine samples from control group and four time points in EAU group (days 5, 8, 12, and 16) were analyzed by label-free DIA workflow.

To generate spectral library A, fractions separated by spin column were analyzed by DDA-MS, and then were processed using proteome discoverer (version 2.3) and Spectronaut Pulsar X. The library A included ten DDA analyses of fractions resulting 1255 protein groups and 6879 peptides with at least one unique peptides and Q value<0.01. DIA-MS raw data files acquired with 26 refined isolation windows from the forty individual urine samples were loaded into Spectronaut Pulsar X using the library A. Overall, a total of 704, and in average 626, protein groups were identified with forty biological replicates. All identification and quantitation details are listed in supporting Table S3.

One hundred proteins were significantly changed in EAU urine samples compared to control ones (1.5-fold change, P<0.05). The overlap of differential expressed proteins identified at different EAU stages is shown by a Venn diagram (Fig. 3). There were 37, 33, 37, and 44 changed urinary proteins on days 5, 8, 12, and 16, respectively, after bovine IRBP immunization (Table S4-7).

**Figure 3.**
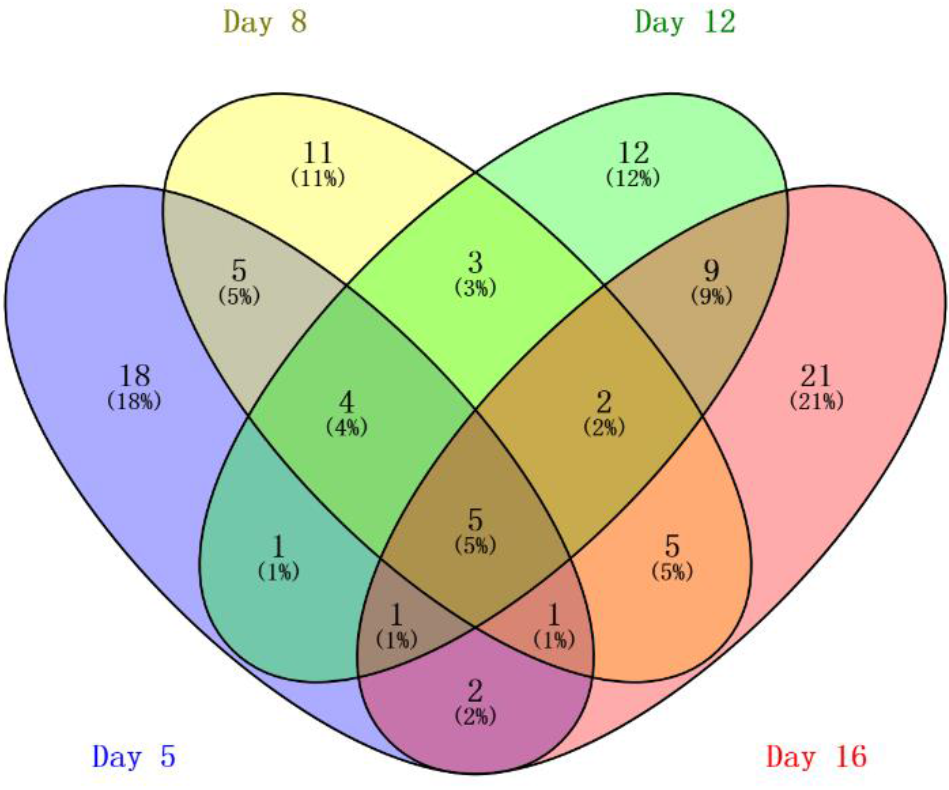
The Vein diagram of the differentially expressed urinary proteins on day 5, 8, 12 and 16, respectively.

Fifteen urinary proteins changed significantly both on day 5 and 8, when there is no clinical manifestations and histopathological ocular damages (Table 1). Suggesting the potential for these urinary proteins to be used for the early detection of uveitis. Among these differential expressed proteins, 9 proteins consistently changed on day 12 when the EAU severity peaked, such as, DEF4, A1AG, NGAL, HPT, ROB1, KNT2, KNT1, MMP8 and KLK9.

**Table 1.** The differentially expressed urinary proteins on early stage of EAU rats

| Uniprot ID | Protein Name | Day 5 |  | Day 8 |  |
| --- | --- | --- | --- | --- | --- |
|  |  | FC | p value | FC | p value |
| Q62714 | Neutrophil antibiotic peptide NP-4 (DEF4) | 4.3 | 6E-06 | 3.0 | 7E-05 |
| P02764 | Alpha-1-acid glycoprotein (A1AG) | 3.9 | 2E-05 | 2.0 | 3E-09 |
| P30152 | Neutrophil gelatinase-associated lipocalin (NGAL) | 2.9 | 8E-07 | 2.2 | 1E-06 |
| P06866 | Haptoglobin (HPT) | 2.7 | 3E-04 | 1.8 | 3E-06 |
| O55006 | Protein RoBo-1 (ROB1) | 2.5 | 6E-09 | 2.0 | 3E-07 |
| P08932 | T-kininogen 2 (KNT2) | 2.5 | 9E-07 | 1.7 | 7E-06 |
| P01048 | T-kininogen 1 (KNT1) | 2.2 | 3E-06 | 1.7 | 4E-06 |
| O88766 | Neutrophil collagenase (MMP8) | 2.2 | 3E-04 | 2.4 | 3E-04 |
| Q62930 | Complement component C9 (C9) | 2.0 | 4E-05 | 1.6 | 2E-04 |
| Q63515 | C4b-binding protein beta chain (C4BPB) | 1.8 | 4E-03 | 1.6 | 2E-02 |
| P07647 | Submandibular glandular kallikrein-9 (KLK9) | 0.5 | 3E-02 | 0.6 | 5E-02 |
| O54858 | Carboxypeptidase Z (CBPZ) | 0.5 | 2E-03 | 0.5 | 5E-03 |
| Q6IE51 | Serine protease inhibitor Kazal-type 7 (ISK7) | 0.5 | 1E-02 | 0.6 | 8E-03 |
| O54728 | Phospholipase B1, membrane-associated (PLB1) | 0.4 | 5E-02 | 0.4 | 5E-02 |
| P62836 | Ras-related protein Rap-1A (RAP1A) | 0.4 | 9E-03 | 0.5 | 2E-02 |

### 3.3 Gene ontology analysis

The Gene ontology (GO) functional annotation was performed on the differentially expressed proteins in EAU rats identified at the discovery phase. One hundred and two dysregulated proteins at four time points were annotated and classified to be involved with certain biological processes (Fig. 4).

**Figure 4.**
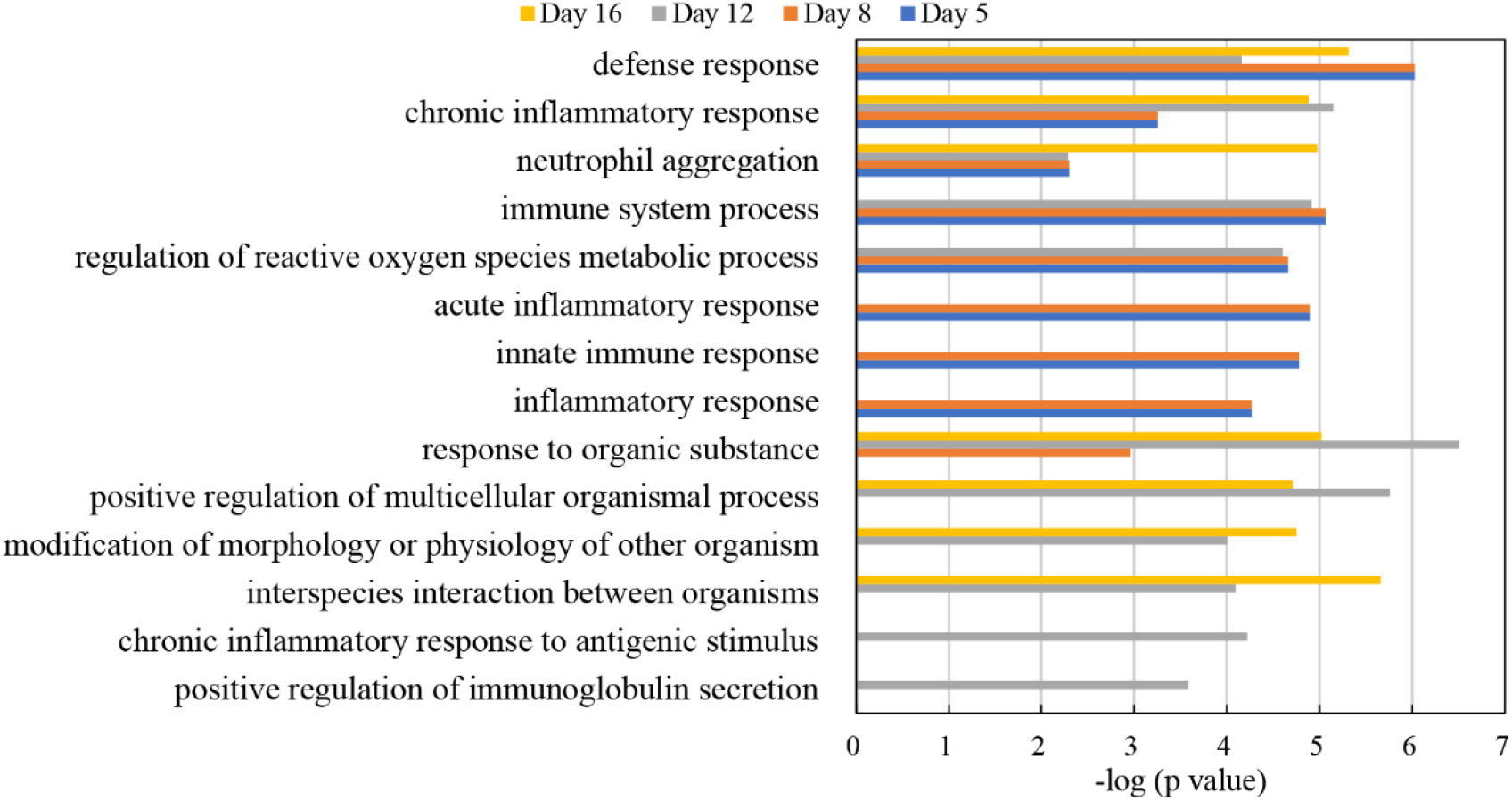
GO enrichment analysis of the differentially expressed proteins at discovery phase in biological processes.

GO enrichment analysis showed that defense response, chronic inflammatory response, neutrophil aggregation and immune system processes were the mainly involved biological processes at four time points. Differential proteins in these GO terms include Protein S100-A8, Neutrophil antibiotic peptide NP-4, Isocitrate dehydrogenase 1 (NADP+), Annexin A2, Ras-related protein Rap-1A, Transforming protein RhoA, Heat shock protein HSP 90-alpha, Carcinoembryonic antigen-related cell adhesion molecule 1 and Glucose-6-phosphate isomerase. The activity of neutrophils is increased and there is an intense neutrophil infiltration in the early stage of inflammation in uveitis [32, 33]. Acute inflammatory response, innate immune response and inflammatory response were significantly enriched at day 5 and 8. Activated innate immunity plays an important role in the pathogenesis of uveitis [34].

### 3.4 PRM validation

At the validation phase, fifty-nine differential expressed proteins were validated in 32 individual urine samples by using parallel reaction monitoring (PRM) method. After further optimization using pooled peptides, forty-four proteins with 150 peptides were finally scheduled for PRM-MS analysis. Overall, seventeen of the forty-four targeted proteins (6 up-regulated and 11 down-regulated) were changed statistically significant at multiple time points (1.5-fold change, p<0.05) (Table 2). The expression trends of these corresponding proteins were consistent with the results from DIA discovery phased. Among these differential proteins, a panel of 12 urinary proteins (MMP8, NGAL, HPT, UROM, RISC, A1AG, TTHY, KNT1, C9, PTER, CBG, and FUCA1) changed significantly even there is no clinical manifestations and histopathological ocular damages in the EAU rats. Four proteins were reported to be associated with uveitis, including MMP8, C9, HTP and CD146.

**Table 2.** The differentially expressed urinary proteins validated by PRM analysis

| Uniprot ID | Protein name | Day 5 |  | Day 8 |  | Day 12 |  |
| --- | --- | --- | --- | --- | --- | --- | --- |
|  |  | FC | p value | FC | p value | FC | p value |
| O88766 | Neutrophil collagenase (MMP8) | 2.2 | 6E-07 | 2.0 | 1E-03 | 2.4 | 2E-04 |
| P30152 | Neutrophil gelatinase-associated lipocalin (NGAL) | 2.1 | 2E-05 | 2.9 | 6E-04 | 3.5 | 3E-04 |
| P06866 | Haptoglobin (HPT) | 2.0 | 1E-04 | 1.6 | 9E-04 | 1.6 | 2E-03 |
| P27590 | Uromodulin (UROM) | 0.6 | 6E-03 | 0.5 | 4E-03 | 0.5 | 2E-02 |
| Q920A6 | Retinoid-inducible serine carboxypeptidase (RISC) | 0.6 | 8E-03 | 0.5 | 1E-03 | 0.4 | 2E-03 |
| P02764 | Alpha-1-acid glycoprotein (A1AG) | 3.0 | 1E-04 | 1.7 | 4E-02 |  |  |
| P02767 | Transthyretin (TTHY) | 0.6 | 4E-03 | 0.6 | 4E-02 |  |  |
| Q62930 | Complement component C9 (C9) | 1.9 | 8E-03 |  |  | 1.5 | 4E-02 |
| P01048 | T-kininogen 1 (KNT1) | 1.7 | 9E-03 |  |  |  |  |
| Q63530 | Phosphotriesterase-related protein (PTER) |  |  | 0.6 | 1E-02 | 0.3 | 2E-03 |
| P31211 | Corticosteroid-binding globulin (CBG) |  |  | 0.7 | 2E-02 |  |  |
| P17164 | Tissue alpha-L-fucosidase (FUCA1) |  |  | 0.6 | 4E-04 |  |  |
| P23785 | Granulins (GRN) |  |  |  |  | 0.6 | 3E-02 |
| P42854 | Regenerating islet-derived protein 3-gamma (REG3G) |  |  |  |  | 0.6 | 6E-03 |
| P45479 | Palmitoyl-protein thioesterase 1 (PPT1) |  |  |  |  | 0.4 | 3E-02 |
| Q6DGG1 | Protein ABHD14B (ABHEB) |  |  |  |  | 0.5 | 1E-02 |
| Q9EPF2 | Cell surface glycoprotein MUC18 (CD146) |  |  |  |  | 0.6 | 3E-02 |

Matrix metalloproteinases (MMPs) comprise a family of zinc-dependent endopeptidases that function to maintain and remodel tissue architecture. Individual MMPs have been found to be expressed in all eye tissues [35], and the expression of MMPs (MMP2, MMP8, MMP12, and MMP13) is positively associated with increased levels corneal fibrosis and neovascularization [36]. In the aqueous humor of juvenile idiopathic arthritis patients with anterior uveitis, the levels of MMPs (MMP1, MMP3, MMP9) were significantly higher compared to controls [37]. In retinas from EAU rats, the expression of MMPs (MMP7, MMP8, MMP12) were increased compared with naive controls [38]. Besides, MMP8 was also found to be differentially expressed in the urine of autoimmune inflammatory diseases, such as experimental autoimmune myocarditis [39].

Haptoglobin (HPT) and Complement component C9 (C9) were reported to be associated with Behcet’s disease (BD). Uveitis is the most common ocular manifestation in BD, which is a chronic multi-systemic autoimmune inflammatory disease [40]. Previous studies have shown that C9 was significantly increased in plasma of BD patients with relapse [41, 42]. Higher serum HPT levels were obtained in patients with BD compared to controls, and there is no obvious difference between the quiescent and active stages [43, 44]. The plasma level of HPT increased signifcantly in EAU rats compared to controls [13].

Cell surface glycoprotein MUC18 (CD146) is a transmembrane glycoprotein expressed at the junction of endothelial cells[45]. It plays an important role in angiogenesis and vessel remodeling [45, 46]. In age-related macular degeneration patients, serum CD146 level was significantly higher than in the controls [47].

Alpha-L-fucosidase-1 (FUCA1), the enzyme that removes terminal α-L-fucose residues from glycoproteins, is downregulated in some high malignancy cancers [48]. Moreover, FUCA1 was shown to decrease the invasion capability of HuH28 cells by downregulating MMP-2 and MMP-9 [49]. Our results shown the decreased FUCA1 and increased MMP-8.

Previous studies of biomarkers of uveitis were primarily stuck in aqueous humor, blood and tear. In this preliminary study, we first studied urinary protein candidate biomarkers for uveitis using data independent acquisition (DIA) combined with parallel reaction monitoring (PRM) in experimental autoimmune uveitis (EAU) rats. A panel of 12 urinary proteins were identified to be dysregulated on the early stage of uveitis. To further determine the sensitivity and specificity of these dysregulated proteins, urine samples from uveitis patients should be included in further studies. Besides, an in-depth profile of the urine proteome may provide more comprehensive and accurate understanding of the uveitis.

## 4. Conclusion

In this study, twelve candidate urinary biomarkers of uveitis were uncovered at the early stage of EAU rats, when there is no clinical manifestations and histopathological ocular damages. Our results suggest that the urinary proteome might reflect the pathophysiological changes in EAU. These dysregulated proteins were potential biomarkers for early diagnosis of uveitis.

## Acknowledgement

The authors declare that they have no competing interests. National Key Research and Development Program of China (2018YFC0910202, 2016YFC1306300); Beijing Natural Science Foundation (7172076); Beijing cooperative construction project (110651103); Beijing Normal University (11100704); Peking Union Medical College Hospital (2016-2.27).

---

**Figure S2:**
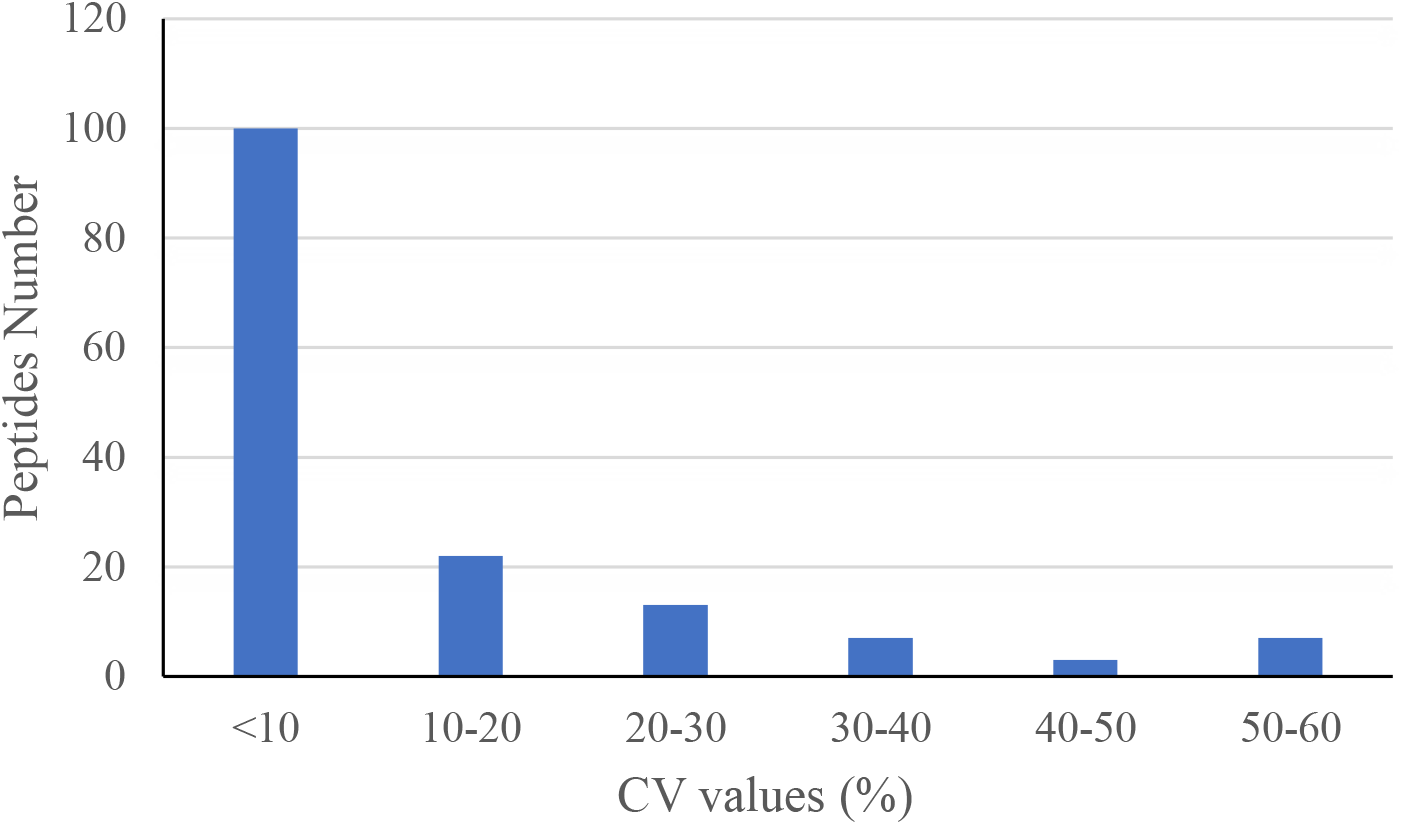
CV values of the 150 PRM targeted peptides.

